# FBL as a carrier mediate H2Aub degradation in the lysosome-like structures

**DOI:** 10.1101/2025.06.30.662258

**Authors:** Jinli Jin, Hanghang Lou, Xuxiang Chen, Senyu Ma, Yunxiao Li, Junli Zhang, Lu Yang

## Abstract

Under glucose starvation, mammalian cells form lysosome-like structures within the nucleoli to mediate the metabolic shift of carbon sources from glucose to amino acids generated through H2Aub degradation for energy supply, but the mechanism by which lysosome-like structures mediate the degradation of monoubiquitinated histones during glucose starvation remains unclear. Through screening, we discovered that the fibrillarin (FBL) protein in the nucleolus acts as a carrier for mono-ubquitinated histone H2A (H2Aub), regulating the balance between BMI1-mediated mono-ubiquitination of H2A and USP36-mediated deubiquitination of H2Aub. Under glucose starvation, FBL undergoes aggregation, leading to a significant reduction in its interaction with BMI1, accompanied by markedly decreased binding to histone H2A and H2Aub.

Concurrently, Midnolin, which serves as a receptor for monoubiquitinated histones, exhibits increased interaction with FBL during glucose starvation. USP36, FBL, Midnolin, and BMI1 collectively form a complex responsible for degrading monoubiquitinated histones. Furthermore, knockdown of FBL, BMI1, or USP36 resulted in cell cycle arrest at S phase and severely compromised cell viability. In summary, we identified a lysosome-like structure in the nucleolus that mediates the degradation of H2Aub. This structure plays a critical role in regulating cell cycle progression and survival, and may offer novel potential therapeutic targets for cancer.

## Introduction

Fibrillarin (FBL) is highly evolutionarily conserved and primarily localizes to the nucleolus and Cajal bodies within the nucleolus [Ref]^1,2^. Its core functions include rRNA methylation, RNA processing, and ribosome assembly [Ref]^3-5^. Additionally, FBL has been shown to mediate methylation of histone H2A in yeast and human cells. For example, in both yeast and human cells, FBL primarily mediates methylation of histone H2A at position 104 (Q105) [Ref]^6,7^. This FBL-dependent H2A methylation is confined to the nucleolus^7^ [Ref], but why this H2A methylation only found in nucleolus still to be elucidated. Studies report that SIRT7-dependent deacetylation regulates FBL’s activity, thereby modulating H2A methylation levels, which in turn influence rRNA synthesis and cell cycle progression^8^ [Ref]. Furthermore, aberrant expression of FBL is associated with various diseases, including cancer, autoimmune disorders, and viral infections^9-11^ [Ref]. However, whether the interaction between FBL and H2A contributes to these disease pathologies remains unclear.

BMI1 and RING1B in the PRC1 complex form a heterodimer, which serves as the primary E3 enzyme responsible for mono-ubiquitination of H2Aub (H2A mono-ubiquitination) in cells ^12^[Ref]. Notably, knockdown of BMI1, but not RING1B, results in a significant reduction in H2Aub levels^13,14^ [Ref]. Bmi1 is overexpressed in various cancers, where it promotes tumor initiation and progression by regulating the cell cycle, enhancing proliferation, and inhibiting apoptosis^15,16^ [Ref]. For instance, in liver cancer, pancreatic cancer, and neuroblastoma, Bmi1 exerts its effects through mechanisms such as modulating the Ink4a/Arf pathway and silencing Hox genes^17,18^ [Ref]. While glucose starvation (GS) has been shown to reduce H2Aub levels, how this process may contribute to tumor cell survival still need to be clarified^19^ [Ref].

USP36 is a nucleolus-localized E3 deubiquitinating enzyme^20,21^ [Ref] that exhibits conserved activity in removing mono-ubiquitination from H2Bub (H2B ubiquitination) *in vitro*^22^ [Ref]. However, whether it catalyzes the deubiquitination of H2Aub in cellular contexts and its physiological functions remain to be elucidated. Midnolin has been reported as a nuclear receptor for ubiquitinated proteins, binding to nuclear ubiquitinated substrates and facilitating their degradation via nuclear proteasomes^23,24^ [Ref]. Furthermore, our previous study has revealed that Midnolin acts as a receptor for H2Aub, mediating its degradation in nucleolar lysosomes during glucose starvation.

In this study, we elucidated the mechanism by which lysosome-like structures in the nucleolus degrade monoubiquitinated histone H2Aub. Through screening, we identified FBL exhibits strong colocalization with lysosome-like structures (LLS) in the nucleolus. Immunoprecipitation-mass spectrometry (IP-MS) experiments further revealed that FBL interacts with BMI1 and USP36, suggesting its role as a carrier for H2Aub to balance histone H2A mono-ubiquitination and H2Aub deubiquitination. Notably, under glucose starvation, FBL displayed enhanced colocalization with Midnolin in the nucleolus. Collectively, our results suggest the structure and mechanism of the lysosome-like pathway in the nucleolus, which degrade H2Aub into amino acids and regulate the availability of amino acids in cell cycle regulation.

## Results

### FBL acts as a carrier for H2Aub to mediate its degradation

Our previous study identified the presence of Lamp1 in the nucleolus, which participate in the degradation of H2Aub, yet the precise degradation mechanism remains unclear. To investigate how ubiquitinated H2A (H2Aub) is degraded within the nucleolus, we first screened nucleolar proteins to identify candidates colocalizing with lysosomal marker Lamp1 (Data not shown). Intriguingly, we observed strong colocalization between FBL and Lamp1 across multiple cell lines and under various conditions, with FBL encircling LAMP1 structures (Fig. 1a). Moreover, combined treatment with glucose starvation and chloroquine (CQ) induced the formation of large foci of FBL and Lamp1 around the nucleolus (Fig. 1a).

**Figure 1.**
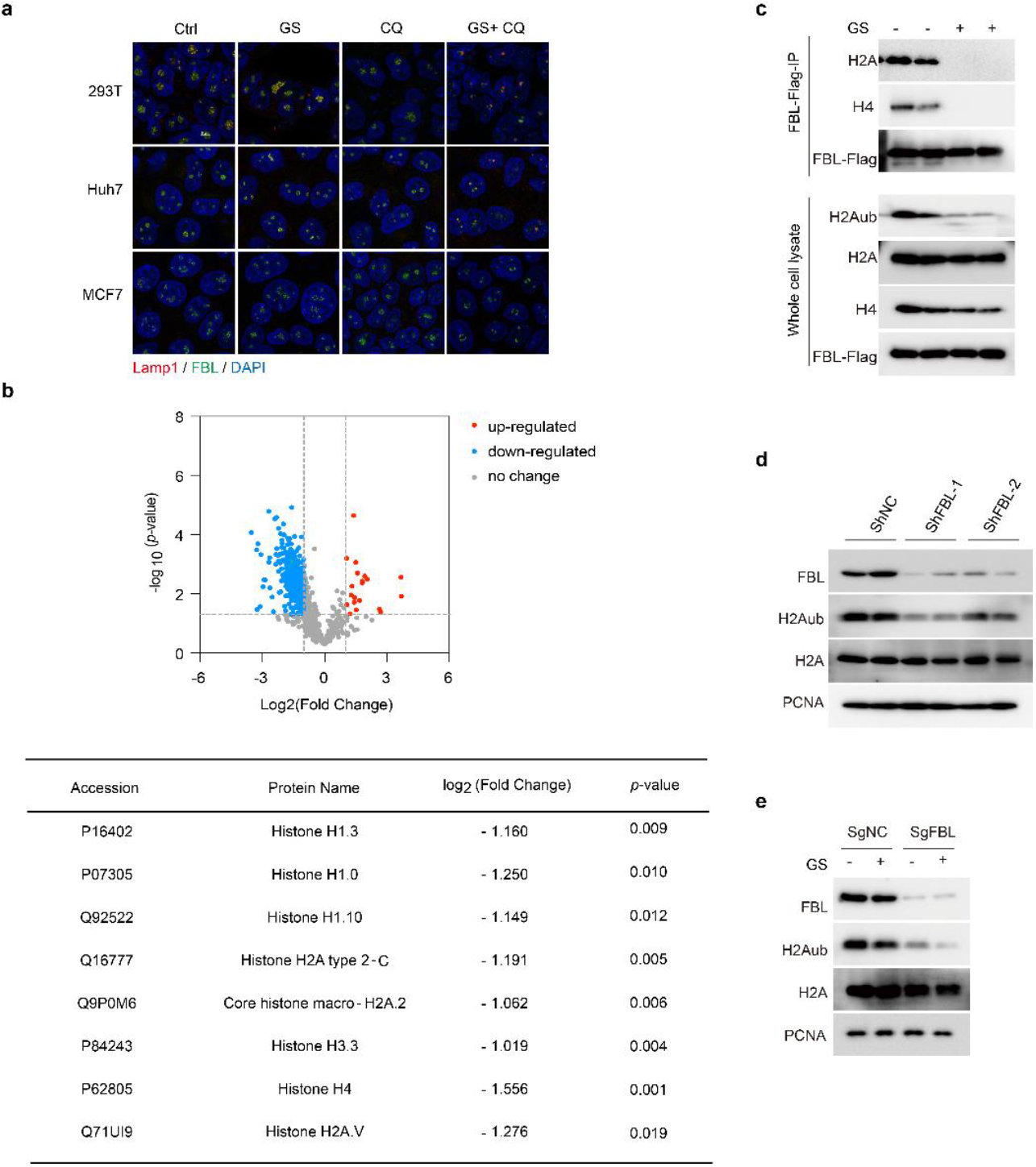
FBL is involved in the intranucleolar lysosome-related pathway a.Combined treatment with glucose starvation and CQ induces the formation of large aggregates containing FBL and the lysosomal marker LAMP1, with persistent co-localization between FBL and LAMP1. b.MS data reveal that GS treatment significantly reduces histone levels bound to FBL. c.WB analysis demonstrates a marked decrease in FBL-associated histone levels upon GS treatment. d.FBL knockdown by shRNA leads to reduced H2Aub levels. e.Glucose starvation further diminishes H2Aub levels in FBL-knockdown cells.

To investigate the role of FBL in the degradation of H2Aub, we employed immunoprecipitation-mass spectrometry (IP-MS) to analyze how glucose starvation affects FBL-interacting proteins. We found that glucose starvation significantly reduced the association of histones with FBL (Fig. 1b, Supplementary Table1). Further validation using Western blotting confirmed that glucose starvation markedly decreased the levels of histones and H2Aub bound to FBL (Fig. 1c). This observation aligns with previous reports demonstrating an interaction between FBL and H2A. our earlier results shown that nucleolar histones are localized around Lamp1, suggesting FBL may act as a carrier for H2A.

Knockdown of FBL by shRNA and sgRNA consistently reduced H2Aub levels (Fig. 1d, e), and glucose starvation further amplified this reduction (Fig. 1e). Through mass spectrometry, immunofluorescence, and Western blotting analyses, we demonstrated that FBL serves as a carrier in the degradation process of H2Aub.

### FBL acetylation was involved in H2Aub degradation

Previous studies have indicated that FBL acetylation plays a role in FBL phase separation. To explore whether FBL acetylation promotes glucose starvation-induced degradation of H2Aub, we knocked down SIRT7, the primary deacetylase responsible for FBL. We found that SIRT7 knockdown by siRNA and shRNA enhanced the glucose starvation-induced reduction of H2Aub levels (Fig. 2a, b), suggesting that FBL acetylation contributes to the degradation of H2Aub under glucose starvation.

**Figure 2.**
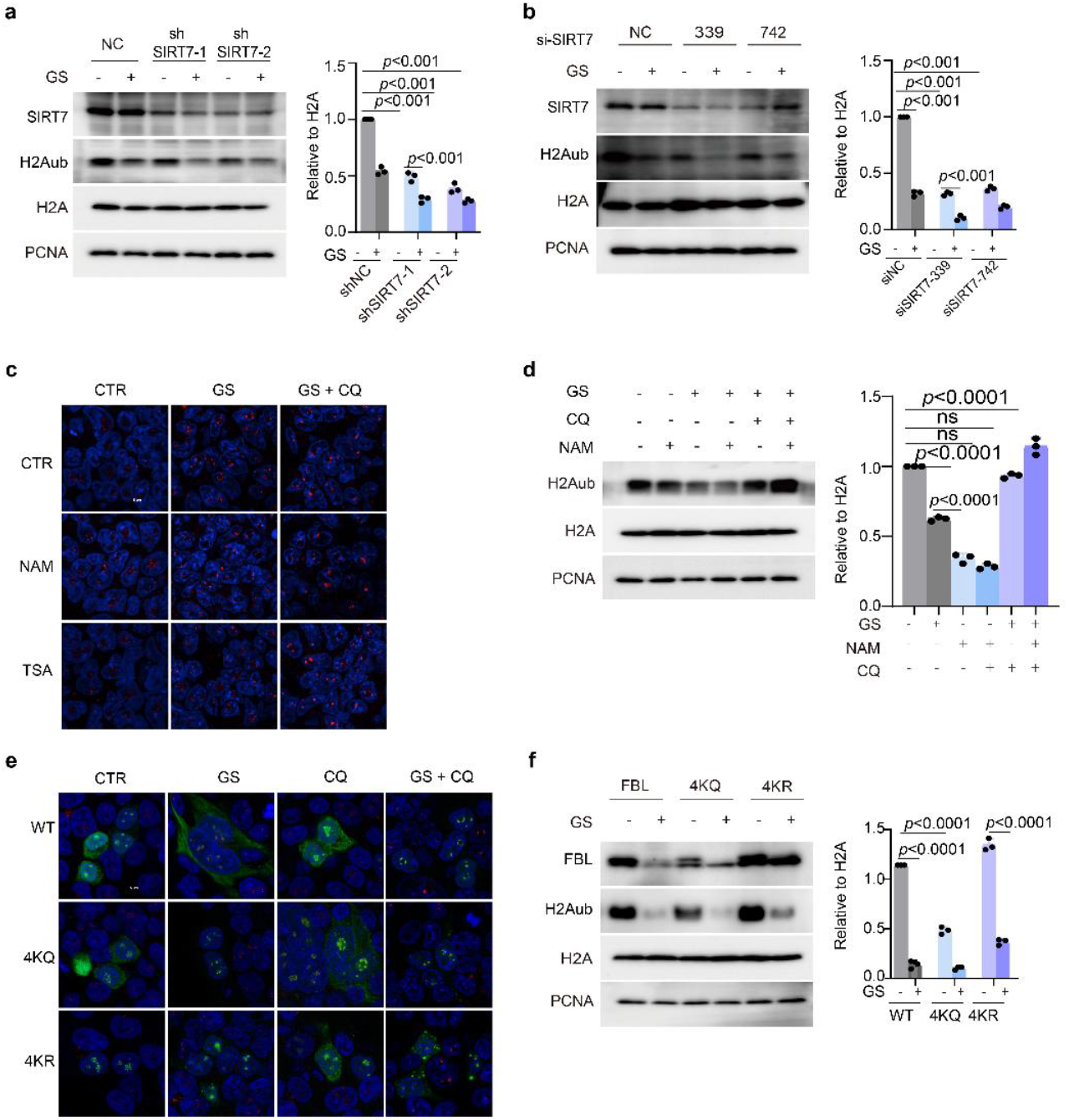
FBL acetylation regulates the intranucleolar lysosome-related degradation pathway. Western blot analysis and quantification show that the acetylation inhibitor NAM enhances glucose starvation-induced LAMP1 aggregation. Western blot analysis and quantification show that the acetylation inhibitor NAM exacerbates glucose starvation-induced reduction of H2Aub levels. The acetylation inhibitor TSA significantly increases H2Aub levels (SOM). SIRT7 knockout exacerbates glucose starvation-induced reduction of H2Aub levels. Western blot analysis and quantification show that reconstitution of FBL-knockout (KO) cells with plasmids expressing differentially acetylated FBL variants to investigate the relationship between FBL acetylation, aggregation propensity, and glucose starvation response. FBL-Flag were transfected into 293T. Then cells were treated with glucose starvation. Endogenous Lamp1(red) and overexpressed FBL-Flag (green) were stained with rabbit anti-Lamp1 antibody and mouse anti-Flag antibody respectively, and were imaged by confocal microscopy. Western blot analysis and quantification show that reconstitution of FBL-KO cells with FBL plasmids reveals coordinated changes in H2Aub levels and FBL content across variants.

We then employed the SIRT7 inhibitor NAM to enhance FBL acetylation. We observed that NAM further exacerbated the glucose starvation (GS)-induced increase in lysosome-like aggregate (LLA) formation (Fig. 2c), whereas Histone deacetylase inhibitor Trichostatin A (TSA) showed no significant promotive effect on FBL aggregation. Furthermore, consistent with SIRT7 knockdown effects, NAM augmented GS-triggered degradation of ubiquitylated H2A (H2Aub) (Fig. 2d), while TSA conversely attenuated H2Aub reduction (Data not shown). These results demonstrate that FBL acetylation mediates H2Aub degradation under glucose starvation, whereas histone acetylation itself suppresses degradation of histones via the H2Aub pathway (Fig. 2a).

Upon overexpressing distinct acetylated variants of FBL in cells, we observed that glucose deprivation (GS) treatment markedly enhanced foci formation of the 4KQ mutant (acetylation-mimetic at all four lysine residues) (Fig. 2d). In contrast, the 4KR mutant (deacetylation-mimetic at all four lysine residues) suppressed FBL foci formation, implying that FBL acetylation regulates glucose starvation-induced ubiquitylated H2A (H2Aub) degradation (Fig. 2d). In FBL-knockdown cells reconstituted with either 4KQ or 4KR mutants, the 4KQ variant accelerated the degradation of both FBL and H2Aub, whereas 4KR attenuated their clearance (Fig. 2e). Collectively, these findings establish that FBL acetylation modulates glucose-dependent H2Aub degradation.

### FBL serves as a carrier that mediates the dynamic ubiquitination and deubiquitination of H2Aub

Analysis of mass spectrometry data revealed that glucose starvation concomitantly reduced FBL interaction with RING1B (a Polycomb repressive complex 1 subunit) and BMI1 (a Polycomb RING finger protein) (Fig. 3a). This reduction in RING1B-FBL interaction was further validated by immunoblotting (Fig. 3b). Strikingly, BMI1 knockdown partially rescued the glucose starvation-induced decrease in H2Aub, which mechanistically supports that glucose deprivation promotes H2Aub degradation by attenuating interactions between histones (H2A) and their mono-ubiquitination machinery (BMI1) (Fig. 3c).

**Figure 3.**
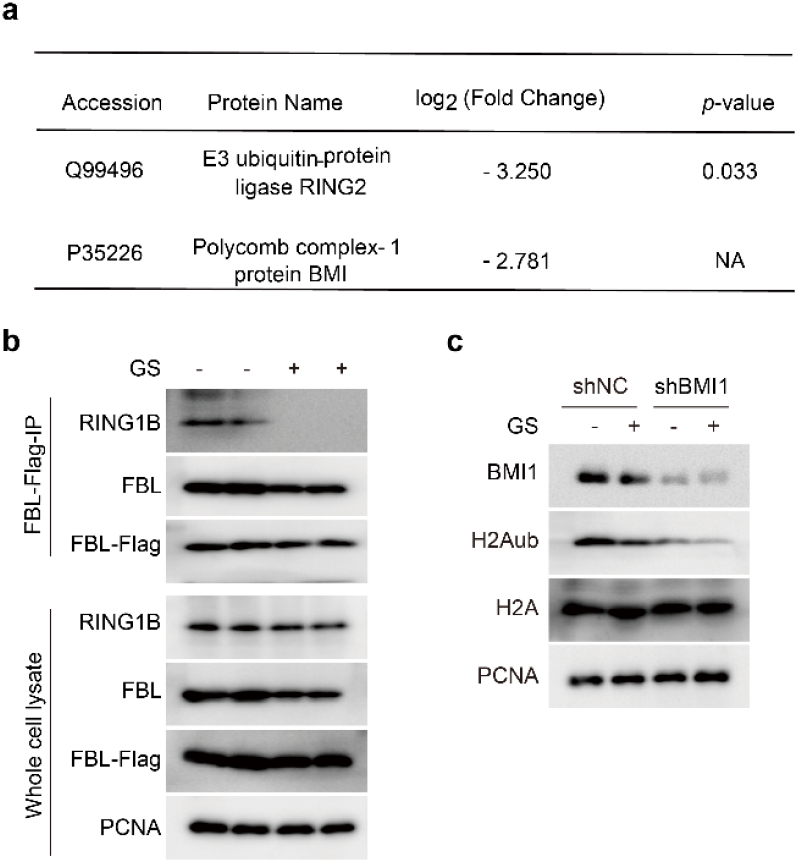
FBL interacts with Bmi1 and RING1B to facilitate monoubiquitination of histone H2A a.MS analysis shows glucose starvation significantly reduces FBL, RING1B, and Bmi1 levels. b.Western blot (WB) analysis demonstrates glucose starvation markedly diminishes the FBL-RING1B interaction. c.Bmi1 knockdown suppresses the glucose starvation-induced reduction of H2A ubiquitination (H2Aub).

### GS treatment mediates the reduction of H2Aub levels through USP36

Further analysis of FBL-interacting proteins revealed an association between USP36 and FBL (Fig. 4a). USP36, previously reported to localize in the nucleolus and mediate *in vitro* deubiquitination of monoubiquitinated H2B (H2Bub), was confirmed to interact with FBL through FBL immunoprecipitation (IP) assays (Fig. 4b). Immunofluorescence assays demonstrated USP36 encircling FBL within the nucleolus under both normal nutrient conditions and glucose starvation (Fig. 4c). Overexpression of USP36 in cells significantly reduced H2Aub levels (Fig.4d), whereas knockdown of USP36 inhibited the glucose starvation-induced reduction in H2Aub levels (Fig. 4e). Collectively, these findings demonstrate that USP36 mediates the degradation of monoubiquitylated histone H2A (H2Aub) within nucleoli.

**Figure 4.**
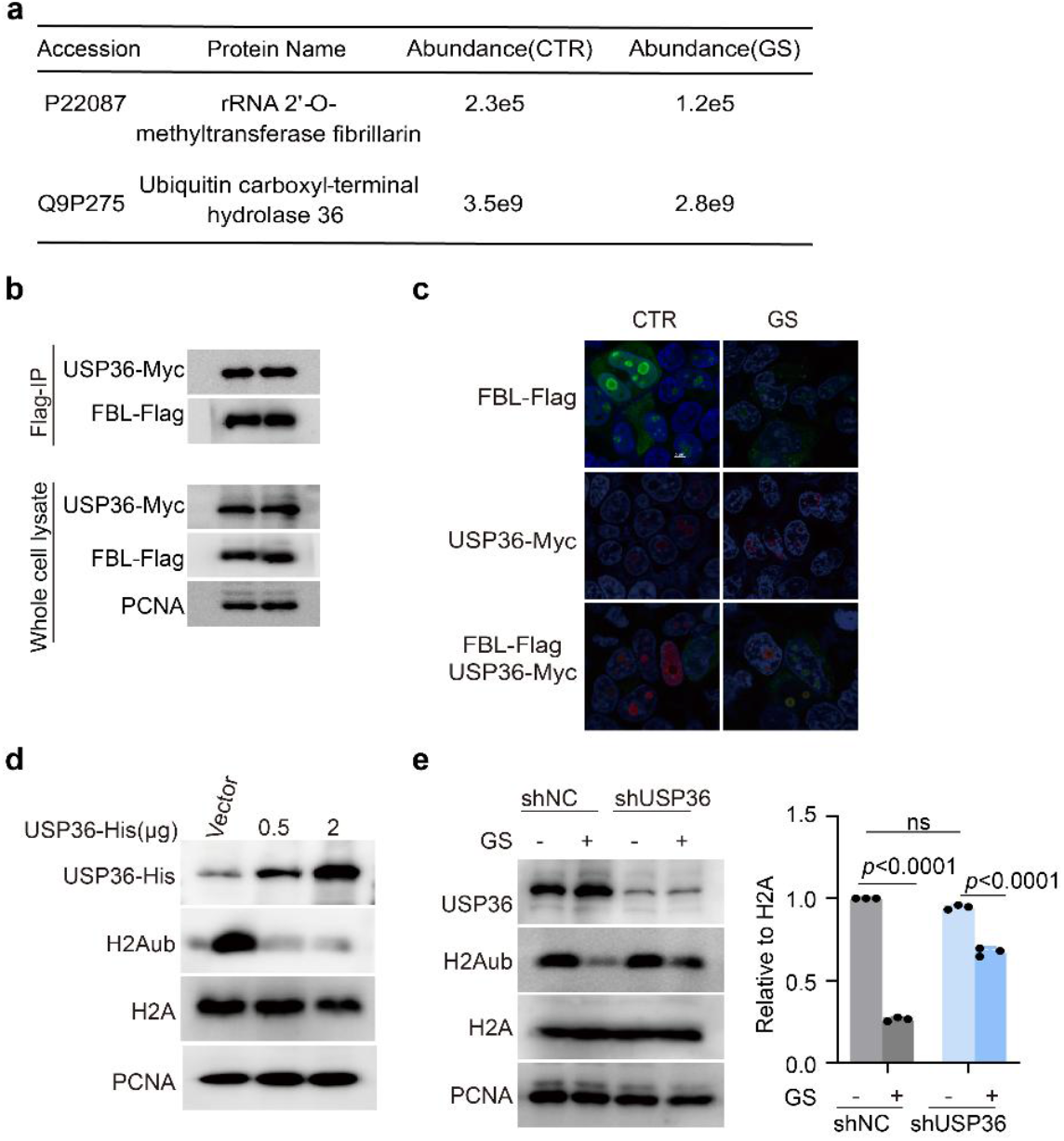
FBL promotes H2A ubiquitination (H2Aub) deubiquitination through its interaction with USP36. FBL promotes H2A ubiquitination (H2Aub) deubiquitination through its interaction with USP36. a. Mass spectrometry analysis demonstrates the interaction between FBL and USP36. Western blotting with immunoprecipitation (WB-IP) confirms the FBL-USP36 interaction. Immunofluorescence reveals USP36 encircling FBL within the nucleolus. FBL-flag, USP36-myc were transfected into 293T separately or at the same time. Then cells were treated with glucose starvation. Overexpressed FBL-Flag (green) and USP36-Myc (red) were stained with mouse anti-Flag and rabbit anti-Myc antibody respectively, and were imaged by confocal microscopy. Overexpression of USP36 reduces cellular H2A ubiquitination (H2Aub) levels. Western blot analysis and quantification show that USP36 knockdown attenuates glucose starvation-induced H2Aub reduction.

### Lysosome-like structures in the nucleolus

Previous studies suggested that Midnolin may act as a receptor mediating the degradation of H2Aub in the nucleolus, while this study reveals that FBL likely serves as a carrier for H2Aub to facilitate its nucleolar degradation. Consistent with this, Immunofluorescence assays demonstrated that glucose starvation increases the interaction between Midnolin and FBL (Fig. 5a). Western blot analysis further demonstrated that glucose starvation significantly enhances the interaction between FBL and Midnolin (Fig. 5b). Furthermore, glucose starvation also increased USP36 and Midnolin colocalize within the nucleolus (Fig. 5c, d). Collectively, we propose a lysosome-like structure in the nucleolus (Fig. 5e): FBL acts as a carrier for H2Aub, mediating both its mono-ubiquitination and deubiquitination to maintain dynamic equilibrium of H2Aub modification levels. Under glucose starvation, the reduction in FBL-bound monoubiquitinated histones diminishes, while increase in Midnolin-H2Aub interactions, thereby promoting H2Aub entry into the nucleolar lysosome-like structure for degradation.

**Figure 5.**
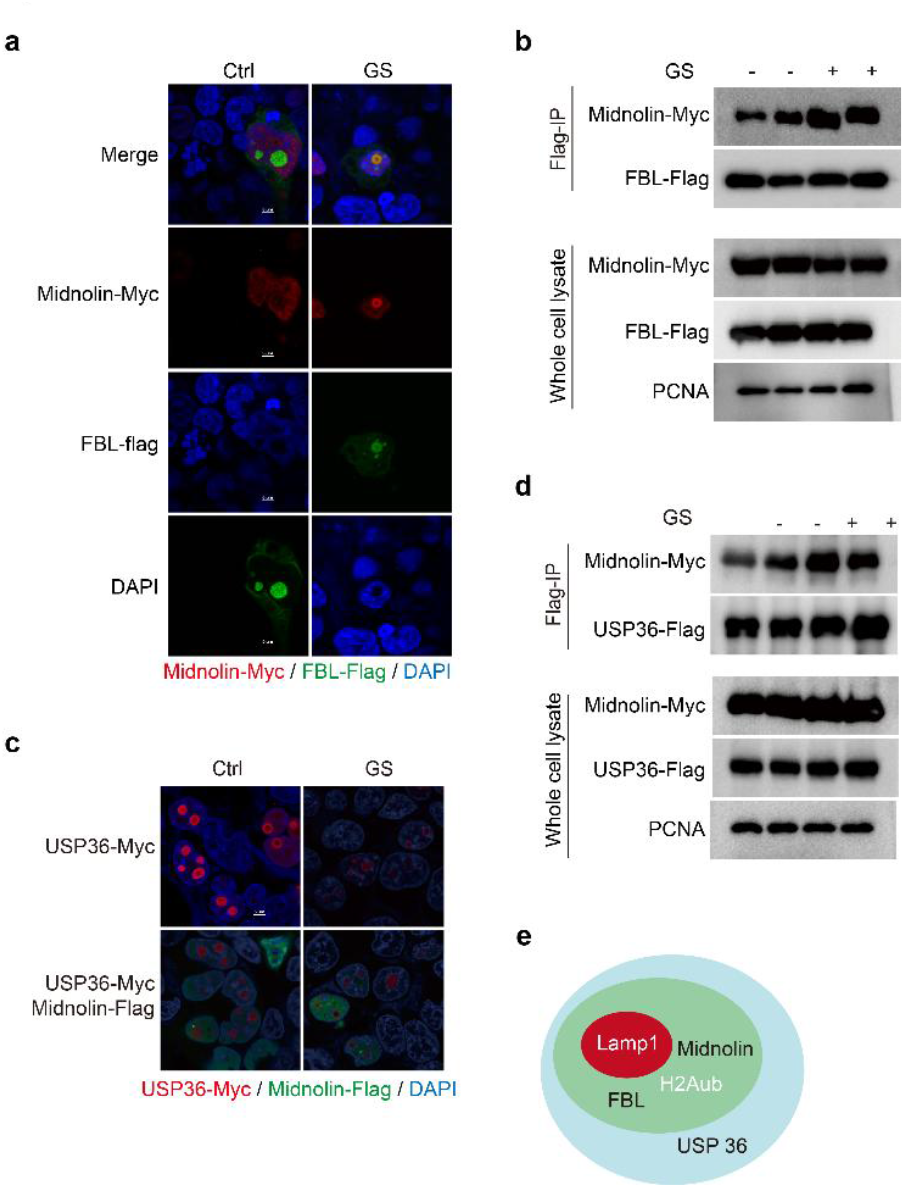
Glucose starvation enhances the interaction between FBL and Midnolin. f.Immunofluorescence demonstrates enhanced FBL-Midnolin interaction upon glucose starvation. Midnolin-Myc and FBL-flag were transfected at the same time. Overexpressed FBL-Flag (green) and Midnolin-Myc (red) were stained with mouse anti-Flag and rabbit anti-Myc antibody respectively, and were imaged by confocal microscopy. a. Western blot with immunoprecipitation confirms increased FBL-Midnolin binding under glucose-deprived conditions. g. Immunofluorescence analysis reveals strengthened USP36-Midnolin co-localization in response to glucose starvation. USP36-Myc and Midnolin-flag were transfected at the same time. Overexpressed Midnolin-Flag (green) and USP36-Myc (red) were stained with mouse anti-Flag and rabbit anti-Myc antibody respectively, and were imaged by confocal microscopy. b.WB-IP assays verify augmented USP36-Midnolin interaction following glucose starvation treatment. c.FBL, BMI1, USP36, Midnolin and H2Aub form a Lysosome-like structure in the nucleolus which may contribute to the degradation of H2Aub.

### FBL affects the cell cycle through its role in mediating H2Aub degradation

Our previous study has demonstrated that glucose starvation induces cell cycle arrest. During the generation of knockdown cell lines, we observed that cells with shFBL, shBMI1, or shUSP36 exhibited severely compromised viability, with USP36-depleted cells showing the most pronounced deterioration (**Data not shown**). This suggests that USP36-mediated deubiquitination of monoubiquitylated histones plays a critical role in cellular survival. To investigate how proteolysis-dependent energy mobilization regulates cell cycle progression, we analyzed cell cycle profiles following FBL, BMI1 or USP36 depletion. FBL, BMI1 and USP36-deficient cells displayed significant S-phase arrest (Fig. 6a-d), which is consistent with the hypothesis that their roles in degradation of H2Aub to provide amino acids.

**Figure 6.**
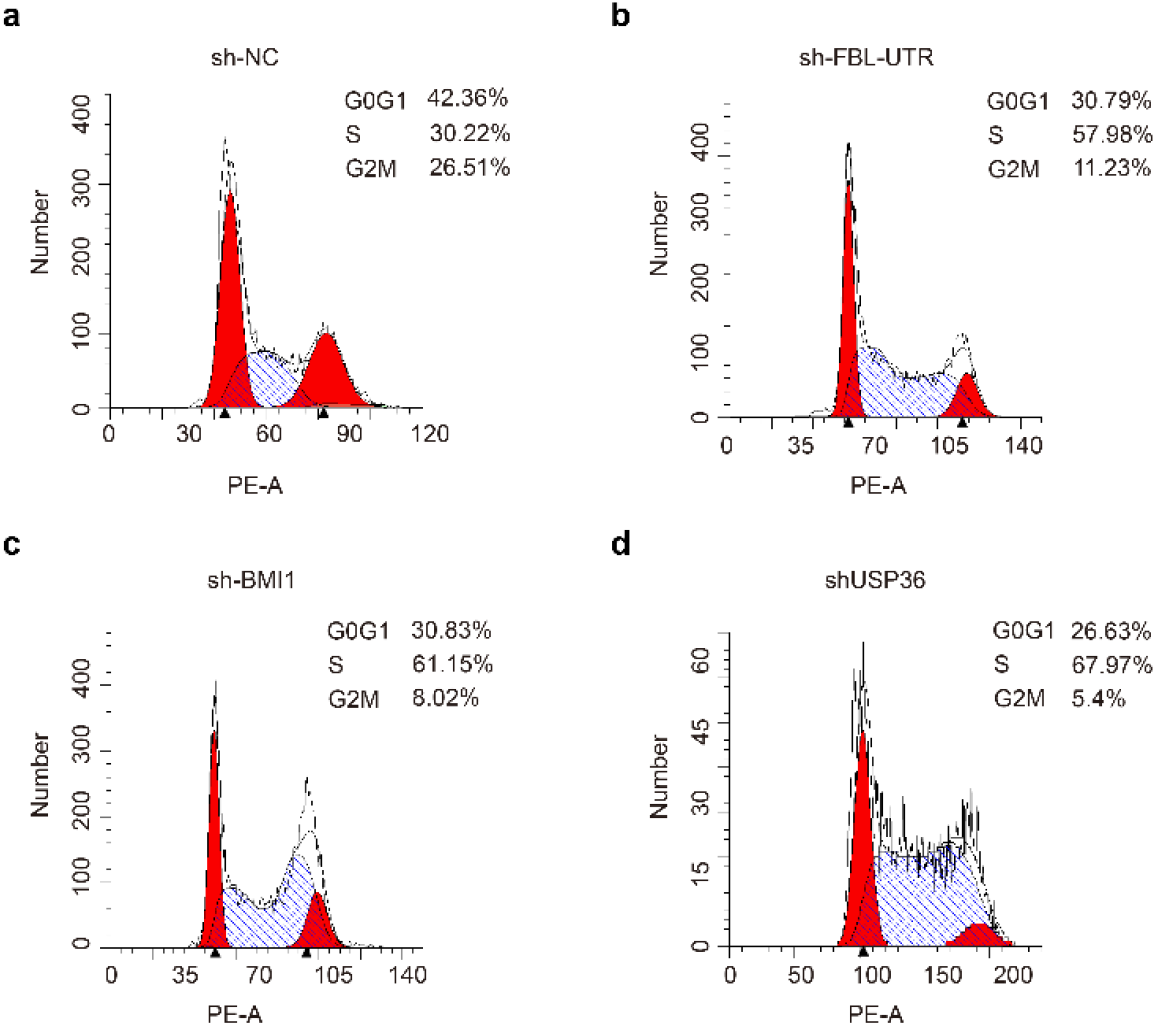
Knockdown of FBL, Bmi1, and USP36 suppresses cell cycle progression into S phase. a.shNC is a reference for cell cycle analysis. b.shFBL induces S-phase arrest in cells. c.shBMI1 triggers cell cycle arrest at S phase. d.shUSP36 results in S-phase accumulation of cells.

## Discussion

Our previous studies revealed the existence of lysosome-like structures within nucleoli that may play significant roles during the carbon source transition from glucose to amino acids. In the current investigation, we explored the specific mechanisms by which these lysosome-like structures in the nucleus degrade monoubiquitinated histone H2Aub. We identified FBL as a critical transporter of H2Aub that regulates its homeostasis through dual molecular interactions: (1) facilitating H2A monoubiquitination via binding with BMI1, and (2) mediating H2Aub deubiquitination through association with USP36. Furthermore, we discovered that Midnolin serves as a receptor for monoubiquitinated histone protein H2Aub. Under glucose starvation conditions, enhanced interaction between Midnolin and FBL promotes the degradation of monoubiquitinated H2Aub. This coordinated mechanism maintains dynamic equilibrium of H2Aub levels through precise regulation of ubiquitination-deubiquitination cycles and nutrient-responsive degradation.

FBL is a highly conserved nucleolar protein found across species ranging from archaea to humans^25,26^ [Ref]. FBL catalyzes S-adenosylmethionine (SAM)-dependent methylation reactions targeting both rRNA and histone H2A^6,27^ [Ref]. Our study demonstrates that FBL interacts with BMI1 and RING1B subunits of the PRC1 complex to promote H2A mono-ubiquitination. Notably, glucose starvation substantially attenuates FBL’s interaction with both H2A and H2Aub, Previous studies have established that glucose deprivation activates AMPK^28^ [Ref], and activated AMPK triggers nucleolar export of SIRT7 ^29^[Ref]. SIRT7 knockdown increases FBL acetylation while reducing its binding to H2A^8^ [Ref], consistent with our observation of diminished FBL-H2A interaction under glucose starvation. Beyond acetylation, whether glucose deprivation affects FBL’s interaction with H2A through additional post-translational modifications, such as phosphorylation, requires further investigation. Furthermore, since FBL-mediated H2A methylation appears restricted to the nucleolus, the potential role of H2A methylation in regulating nucleolar H2Aub levels merits exploration. Clinically, FBL is overexpressed in multiple malignancies including breast, prostate, and liver cancers, correlating with enhanced proliferation, drug resistance, and poor prognosis^11,30,31^ [Ref]. FBL knockdown suppresses tumor growth^32^ [Ref], suggesting that targeted modulation of FBL-H2Aub interactions might represent a promising therapeutic strategy for cancer intervention.

Recent studies have revealed that PRC1 not only mediates H2A mono-ubiquitination on nucleosomes but also facilitates this modification through liquid-liquid phase separation (LLPS)^33^ [Ref]. In this study, we discovered that within the LLPS-enriched nucleolar environment, BMI1 and RING1B, in complex with FBL, catalyze H2A mono-ubiquitination, a process critically involved in glucose starvation-induced H2Aub degradation. Investigating how BMI1 modulates FBL-H2A interactions provides novel insights into the context-dependent physiological roles of BMI1. For instance, BMI1 plays essential roles in hematopoietic stem cells, neural stem cells, and various adult stem cells by repressing tumor suppressor genes such as p16, Ink4a, and p19Arf to sustain stem cell self-renewal capacity^18,34,35^ [Ref]. Moreover, BMI1 is overexpressed in numerous cancers, where it drives tumorigenesis and progression by modulating cell cycle progression, promoting proliferation, and inhibiting apoptosis^36-38^ [Ref]. For example, in hepatocellular carcinoma, pancreatic cancer, and neuroblastoma, BMI1 exerts its oncogenic effects through mechanisms including regulation of the Ink4a/Arf pathway and Hox gene silencing ^39^[Ref].

USP36 (Ubiquitin-Specific Protease 36) is a deubiquitinating enzyme that predominantly functions within the nucleolus^40,41^ [Ref], regulating cellular processes by deubiquitinating and stabilizing multiple critical proteins. USP36 directly interacts with the transcription factor c-Myc, counteracting SCF^(Fbw7)-mediated ubiquitination and degradation of c-Myc through deubiquitination, thereby stabilizing c-Myc protein levels and promoting cancer cell proliferation^42^ [Ref]. Additionally, USP36 contributes to the pathogenesis of diabetic nephropathy by mediating DOCK4 deubiquitination, which activates the Wnt/β-catenin signaling pathway^43^[Ref]. In vitro studies further demonstrate its capacity to catalyze H2Bub deubiquitination across multiple species^22^ [Ref]. Our findings reveal that USP36 maintains a sustained interaction with FBL in the nucleolus and facilitates H2Aub deubiquitination in cellular contexts, suggesting a collaborative role with BMI1 in maintaining dynamic equilibrium of H2Aub levels. The mechanisms by which nucleolar regulation of USP36-FBL interactions modulates H2Aub homeostasis, thereby influencing tumorigenesis and progression, represent a compelling avenue for future investigation.

Tumors are characterized by cells with exceptionally high energy demands^44,45^ [Ref]. Aberrant expression of H2Aub is closely associated with various human diseases, including cancer and neurodevelopmental disorders^46,47^ [Ref]. FBL, BMI1, and USP36 are all critically linked to cell cycle regulation and tumorigenesis^48-50^ [Ref]. In our study, tumor cell viability was severely compromised upon knockdown of FBL, BMI1, or USP36, with USP36 depletion exhibiting more pronounced lethality than FBL or BMI1 knockdown. This hierarchical survival dependency suggests that USP36 and FBL, BMI1 operate within the same pathway, with USP36 positioned upstream of FBL and BMI1. These findings propose a therapeutic strategy to inhibit tumor growth by targeting FBL, BMI1, and USP36 to reduce H2Aub levels, highlighting novel avenues for cancer treatment.

## Methods

### Cell Culture and Transfections

HEK293T cells were cultured in high-glucose DMEM medium (Saivei, G4511-500ML) containing 10% fetal bovine serum (FBS, Novizan, F101) and 1% penicillin-streptomycin (Zhongqiao Xinzhu, CSP006) at 37^°^C with 5% CO_2_. Transfection was performed using Lipofectamine 2000 (Lip2000, Beyotime, BL623B) following the manufacturer’s protocol when the cell confluence reached 50%-60%.

### The product numbers (Catalog numbers) of the antibodies and drugs used are as follows

Ubiquityl-Histone H2A (Lys119) (D27C4) (Cell Sibnsling Technology, 8240S); Ubiquityl-Histone H2B (Lys120) (Cell Sibnsling Technology, 5546S);

K48-linkagePolyubiquitin (D9D5) Rabbit mAb (Cell Sibnsling Technology, 8081S); SirT7 (D3K5A) Rabbit mAb (Cell Sibnsling Technology, 5360S); Bmi1 (Cell Sibnsling Technology, 6964); RING1B (Cell Sibnsling Technology, 5694); Fibrillarin (Abcam, ab5821); Acetyllysine Mouse mAb (PTMBIO, PTM-101); Midnolin Rabbit Polyclonal Antibody (abbiotec, 251274); GFP tag Polyclonal antibody (Proteintech, 50430-2-AP); FLAG tag (Proteintech, 20543-1-AP); USP36 Polyclonal antibody (Proteintech, 14783-1-AP); PCNA Rabbit pAb (ABclonal, A1336); HRP-Labeled Goat Anti-Rabbit IgG (ZSGB-BIO, ZB-2301); Anti-Flag Magnetic Beads (Beyotime, P2115-2ml); Anti-GFP Magnetic Beads (Beyotime, P2132-2ml); Mouse IgG Magnetic Beads (Beyotime, P2171-1ml); Mouse anti-HA antibody (Santa Cruz, sc7392); MYC tag Polyclonal antibody (Proteintech, 16286-1-AP); His-tag Polyclonal antibody (Proteintech, 1001-0-AP); PSMD4 Polyclonal antibody (Proteintech, 14899-1-AP); FLAG mouse (Merck, F1804); LAMP1 Lysosome Marker (Abcam, ab24170); PSME4 Polyclonal antibody (Proteintech, 18799-1-AP); Alexa Fluor 594 Goat anti-Rabbit IgG (Thermo Fisher, A-110037); Alexa Fluor 488 Goat anti-Mouse IgG (Thermo Fisher, A-11029); DAPI solution (Solarbio, C0060). Chloroquine phosphate (MCE, HY-17589); MG132(MCE, HY-13259); TrichostatinA (MCE, HY-15144); N-Acetylmuramic acid-azide (MCE, HY-157573).

### Glucose starvation and drug treatment experiments

Cells were subjected to glucose starvation by washing with phosphate-buffered saline (PBS) and subsequently cultured in glucose-free DMEM (Basai Media, L160KJ) for 0-to 6-hour periods. For MG132 treatment, cells were exposed to 20 uM MG132 (MCE, HY-13259) dissolved in dimethyl sulfoxide (DMSO), while control cells received an equivalent volume of DMSO (final concentration ≤0.1%).

### Immunoprecipitation (IP)

For each experiment, 293T cells were collected from four 10-cm culture dishes. To each dish, 800 uL of ice-cold Western and IP Lysis Buffer (Beyotime, BL509A) was added. Cells were scraped directly into 1.5 mL EP tubes without prior PBS washing and sonicated on ice until the lysate was clarified. The lysates were centrifuged at 12,000 rpm for 10 min, and the supernatants were collected. The supernatants were incubated with rotation at 4°C for 30 min with 10 uL of Mouse IgG Magnetic Beads (Beyotime, 2171-1ml). The beads were then removed using a magnetic stand, and the supernatant was transferred to a new tube containing 20 uL of pre-washed GFP Magnetic Beads (Beyotime, P2132-2ml). This mixture was rotated overnight at 4°C. The next day, the supernatant was removed, and the GFP beads were washed three times with IP Lysis Buffer (5 min each wash). After removing the wash buffer, 50 uL of 1× SDS Loading Buffer (diluted with 1× TBS) was added, vortexed, and boiled for 10 min before being used directly for immunoblotting.

### Western blotting (WB)

293T cells were seeded in 6-well plates. When the cell confluence reached 70%, the control and experimental groups were treated with the corresponding drugs. Cells were lysed on ice for 30 minutes using lysis buffer (containing RIPA buffer, DTT, PMSF, and protease inhibitors). Protein samples were mixed with loading buffer and boiled for 15 minutes.

Protein separation was performed by SDS-PAGE.

### Immunofluorescence (IF)

Cells cultured on coverslips were fixed with 4% paraformaldehyde (Beyotime, P0099-100ml), permeabilized with 0.2% Triton X-100, and blocked with 5% donkey serum (Solarbio, C1052) in PBST. Subsequently, cells were incubated overnight at 4^°^C with the primary antibodies. After washing, cells were incubated with the secondary antibodies concurrently with DAPI staining. Finally, samples were observed and imaged using a Nikon confocal laser scanning microscope.

### SgRNA

The plasmid mixture for transfection was prepared at a mass ratio of Pspax2: Pmd2.G: lentiCRISPR-V2-sgRNA = 1:1:2 for each well of a 6-well plate. Lipofectamine 2000 was used for transfection. After 8 hours, the medium was replaced with fresh DMEM. The supernatant was collected at 48 hours post-transfection, filtered, and used for transduction of new cells (∼50% confluency). Puromycin (Beyotime, ST551) selection was performed continuously at a concentration of 2.5 μg/ml for 48 hours.

### ShRNA

The sense strand sequences of the ShRNA constructs (5’-3’) were synthesized by Gentlegen (Jiangsu) and are as follows:

ShFUTR:5’- ACCGGCGAGAGATGTGTGTTGATATTCTCGAGAATATCAACACACATCTCTCGTTTTTTGAATTC-3’

ShUS36-1:5’- ACCGGGCGGTCAGTCAGGATGCTATTCTCGAGAATAGCATCCTGACTGACCGCTTTTTTGAATTC-3’

ShUS36-2:5’- ACCGGCGTCCGTATATGTCCCAGAATCTCGAGATTCTGGGACATATACGGACGTTTT TTGA-3’

ShMIDNOLIN-1:5’- ACCGGGATGTGAACATCACGTGTTATCTCGAGATAACACGTGATGTTCACATCTTTTTTGAATTC-3’

ShMIDNOLIN2:5’- ACCGGGTCATTATGCACTTTACAGATCTCGAGATCTGTAAAGTGCA-3’

ShSirt7#1: GGGAGUACGUGCGGGUGU

ShSirt7#2: GACGUCAUGCGGCUCCUCA

ShSirt7#3: CAGACACCAUCCUGUGUCU

### SiRNA

si-FBL and si-SirT7 were purchased from Kingbrave Biotech (Wuhan). Transfection was performed using Lipo8000 (Beyotime, catalog number C5033-1ml) with a final siRNA concentration of 50 nM. The culture medium was replaced with fresh medium eight hours post-transfection. Protein extraction was performed for validation within 48-72 hours post-transfection. The sense strand (5’-3’) sequences used in this study were:

siFBL-451: GAUUUCGGAAGGAGAUGACAATT; UUGUCAUCUCCUUCCGAAAUCTT;

siFBL-533: GUGGACCAGAUCCACAUCAAATT; UUUGAUGUGGAUCUGGUCCACTT;

siFBL-852: GACACUUUGUGAUUUCCAUUATT;

UAAUGGAAAUCACAAAGUGUCTT;

siSirT7-339: CGCCAAAUACUUGGUCGUCUATT; AGACGACCAAGUAUUUGGCGTT;

siSirT7-742: CGGGACACCAUUGUGCACUUUTT; AAGUGCACAAUGGUGUCCCGTT;

SirT7-915: GCCGAAGCUUUACAUCGUGAATT; UUCACGAUGUAAAGCUUCGGCTT.

### Cell Cycle Analysis by Flow Cytometry

Cells were harvested and washed with PBS. Trypsin (VivaCell, C3530-0500) was added for digestion, followed by centrifugation at 1000 rpm for 1 minute. Cells were fixed with pre-chilled, ice-cold 70% ethanol at 4^°^C overnight. After centrifugation at 1000 rpm for 5 minutes, cells were washed with PBS and centrifuged again at 1000 rpm for 1 minute. Cells were stained with PI solution containing RNase at 37^°^C in the dark for 30 minutes. Finally, the samples were analyzed by flow cytometry.

### In-gel digestion

Following SDS-PAGE separation of the IP sample protein and Coomassie Brilliant Blue staining, the target protein band was excised and transferred to a 1.5 ml microcentrifuge tube. The gel piece was subjected to reduction and alkylation using TCEP and CAA. After destaining, the gel was digested overnight with trypsin. Peptides were then extracted from the digest using formic acid (FA) and acetonitrile (ACN).

### Load sample

The supernatant was transferred into an injection vial and loaded onto a silica capillary column (75 μm inner diameter, 360 μm outer diameter) using an EASY-nLC™ 1200 system. A 150-min gradient elution program was run at 400 nL/min flow rate: initial 2% Buffer B (0.1% formic acid in 80% acetonitrile) was maintained for 2 min, increased to 8% from 2-8 min, linearly ramped to 25% from 8-118 min, elevated to 40% from 118-150 min, then rapidly raised to 80% within 3 min, followed by a 3-min isocratic phase at 80% concentration. Following nanoelectrospray ionization (NSI), peptide samples were analyzed using an Orbitrap Exploris^™^ 480 mass spectrometer coupled online with ultra-high performance liquid chromatography through a FAIMS Pro™ interface. The FAIMS system operated in dual compensation voltage mode (−45 V and -65 V) with a spray voltage of 2.0 kV applied. Full MS scans were acquired over m/z 350-1500 range at 60,000 resolution (at m/z 200), using an automatic gain control (AGC) target of 300% and maximum injection time of 50 ms. MS/MS spectra were collected at 15,000resolution, with AGC target 75%, 22 ms injection time, and 1-s cycle time using Top Speed acquisition. Precursor ion selection employed a 5×10^4^ intensity threshold, 1.6 m/z isolation window, and 30% normalized collision energy.

### Data search

Mass spectrometric data were processed using Proteome Discoverer^™^ 2.4 software with SEQUEST HT search engine. Tandem mass spectra were searched against the human UniProt database (2019 release) concatenated with its reversed decoy counterpart. Enzymatic digestion parameters were specified as Trypsin/P with up to 2 missed cleavages allowed. Precursor mass tolerance was set to 10 ppm for initial screening and 5 ppm for main search, while fragment ion mass tolerance was fixed at 0.02 Da. Static modifications included cysteine carbamidomethylation, with dynamic modifications of protein N-terminal acetylation and methionine oxidation. The false discovery rate (FDR) was stringently controlled at <1% using target-decoy approach. Proteins identified through untargeted proteomic analysis are compiled in Supplementary Table S1.

## Acknowledgements & Fundings

This research was funded by grants from National Natural Science Foundation of China (32400978) and Start-up Grant for Yellow River Scholar at Henan University. We thank Chensong Zhang for helpful discussion and advice on this project.

## Authors contribution

LY conceived the project and designed the study. LY, JLJ, HHL, XXC, SYM, YXL and JLZ conducted experiments. LY, JLJ, XXC, SYM, HHL, YXL interpreted data. LY, JJL, HHL and XXC contributed to the discussion and critical regents. LY and JJL, HHL wrote the paper.

## Reference

1. Shubina, M. Y., Musinova, Y. R. & Sheval, E. V. Nucleolar methyltransferase fibrillarin: Evolution of structure and functions. Biochemistry (Moscow) 81, 941–950, doi:10.1134/s0006297916090030 (2016).

2. Rodriguez-Corona, U., Sobol, M., Rodriguez-Zapata, L. C., Hozak, P. & Castano, E. Fibrillarin from Archaea to human. Biology of the Cell 107, 159–174, doi:10.1111/boc.201400077 (2015).

3. Elliott, B. A. et al. Modification of messenger RNA by 2′-O-methylation regulates gene expression in vivo. Nature Communications 10, doi:10.1038/s41467-019-11375-7 (2019).

4. Barneche, F., Steinmetz, F. & Echeverriá, M. Fibrillarin Genes Encode Both a Conserved Nucleolar Protein and a Novel Small Nucleolar RNA Involved in Ribosomal RNA Methylation inArabidopsis thaliana. Journal of Biological Chemistry 275, 27212–27220, doi:10.1016/s0021-9258(19)61499-7 (2000).

5. EM Narcisi, C.V.G.M Fechheimer. Fibrillarin, a conserved pre-ribosomal RNA processing protein of Giardia. J Eukaryot Microbiol 45(1):, 105–111, doi:10.1111/j.1550-7408.1998.tb05077.x. (1998).

6. Loza-Muller, L. et al. Fibrillarin methylates H2A in RNA polymerase I trans-active promoters in Brassica oleracea. Frontiers in Plant Science 6, doi:10.3389/fpls.2015.00976 (2015).

7. Tessarz, P. et al. Glutamine methylation in histone H2A is an RNA-polymerase-I-dedicated modification. Nature 505, 564–568, doi:10.1038/nature12819 (2013).

8. Iyer-Bierhoff, A. et al. SIRT7-Dependent Deacetylation of Fibrillarin Controls Histone H2A Methylation and rRNA Synthesis during the Cell Cycle. Cell Reports 25, 2946-2954.e2945, doi:10.1016/j.celrep.2018.11.051 (2018).

9. Tiku, V. et al. Nucleolar fibrillarin is an evolutionarily conserved regulator of bacterial pathogen resistance. Nature Communications 9, doi:10.1038/s41467-018-06051-1 (2018).

10. Li, P. et al. RNA 2’-O-Methyltransferase Fibrillarin Facilitates Virus Entry Into Macrophages Through Inhibiting Type I Interferon Response. Frontiers in Immunology 13, doi:10.3389/fimmu.2022.793582 (2022).

11. Sun, X. et al. FBL promotes cancer cell resistance to DNA damage and BRCA1 transcription via YBX1. EMBO reports 24, doi:10.15252/embr.202256230 (2023).

12. Gray, F. et al. BMI1 regulates PRC1 architecture and activity through homo- and hetero-oligomerization. Nat Commun 7, 13343, doi:10.1038/ncomms13343 (2016).

13. Barbour, H., Daou, S., Hendzel, M. & Affar, E. B. Polycomb group-mediated histone H2A monoubiquitination in epigenome regulation and nuclear processes. Nature Communications 11, doi:10.1038/s41467-020-19722-9 (2020).

14. Jia, L., Zhang, W. & Wang, C.-Y. BMI1 Inhibition Eliminates Residual Cancer Stem Cells after PD1 Blockade and Activates Antitumor Immunity to Prevent Metastasis and Relapse. Cell Stem Cell 27, 238-253.e236, doi:10.1016/j.stem.2020.06.022 (2020).

15. Li, B. et al. Bmi1 drives hepatocarcinogenesis by repressing the TGFβ2/SMAD signalling axis. Oncogene 39, 1063–1079, doi:10.1038/s41388-019-1043-8 (2019).

16. Qin, Z. et al. Co-targeting BMI1 and MYC to eliminate cancer stem cells in squamous cell carcinoma. Cell Reports Medicine 6, doi:10.1016/j.xcrm.2025.102077 (2025).

17. Kim, W. Y. & Sharpless, N. E. The Regulation of INK4/ARF in Cancer and Aging. Cell 127, 265–275, doi:10.1016/j.cell.2006.10.003 (2006).

18. Bruggeman, S. W. M. et al. Bmi1 Controls Tumor Development in an Ink4a/Arf-Independent Manner in a Mouse Model for Glioma. Cancer Cell 12, 328–341, doi:10.1016/j.ccr.2007.08.032 (2007).

19. Zhang, Y. et al. H2A Monoubiquitination Links Glucose Availability to Epigenetic Regulation of the Endoplasmic Reticulum Stress Response and Cancer Cell Death. Cancer Research, doi:10.1158/0008-5472.can-19-3580 (2020).

20. Sun, X.-X. et al. The nucleolar ubiquitin-specific protease USP36 deubiquitinates and stabilizes c-Myc. Proceedings of the National Academy of Sciences 112, 3734–3739, doi:10.1073/pnas.1411713112 (2015).

21. Qin, K. et al. USP36 stabilizes nucleolar Snail1 to promote ribosome biogenesis and cancer cell survival upon ribotoxic stress. Nature Communications 14, doi:10.1038/s41467-023-42257-8 (2023).

22. DeVine, T., Sears, R. C. & Dai, M.-S. The ubiquitin-specific protease USP36 is a conserved histone H2B deubiquitinase. Biochemical and Biophysical Research Communications 495, 2363–2368, doi:10.1016/j.bbrc.2017.12.107 (2018).

23. Peddada, N. et al. Structural insights into the ubiquitin-independent midnolin-proteasome pathway. Proceedings of the National Academy of Sciences 122, doi:10.1073/pnas.2505345122 (2025).

24. Gu, X. et al. The midnolin-proteasome pathway catches proteins for ubiquitination-independent degradation. Science 381, doi:10.1126/science.adh5021 (2023).

25. Kosakovsky Pond, S. L. et al. Fibrillarin evolution through the Tree of Life: Comparative genomics and microsynteny network analyses provide new insights into the evolutionary history of Fibrillarin. PLOS Computational Biology 16, doi:10.1371/journal.pcbi.1008318 (2020).

26. Zhang, X., Li, W., Sun, S. & Liu, Y. Advances in the structure and function of the nucleolar protein fibrillarin. Frontiers in Cell and Developmental Biology 12, doi:10.3389/fcell.2024.1494631 (2024).

27. Ayadi, L., Galvanin, A., Pichot, F., Marchand, V. & Motorin, Y. RNA ribose methylation (2′-O-methylation): Occurrence, biosynthesis and biological functions. Biochimica et Biophysica Acta (BBA) - Gene Regulatory Mechanisms 1862, 253–269, doi:10.1016/j.bbagrm.2018.11.009 (2019).

28. Lin, S.-C. & Hardie, D. G. AMPK: Sensing Glucose as well as Cellular Energy Status. Cell Metabolism 27, 299–313, doi:10.1016/j.cmet.2017.10.009 (2018).

29. Sun, L. et al. Regulation of energy homeostasis by the ubiquitin-independent REGgamma proteasome. Nat Commun 7, 12497, doi:10.1038/ncomms12497 (2016).

30. Yang, L. et al. Phase separation-competent FBL promotes early pre-rRNA processing and translation in acute myeloid leukaemia. Nature Cell Biology 26, 946–961, doi:10.1038/s41556-024-01420-z (2024).

31. Su, H. et al. Elevated snoRNA biogenesis is essential in breast cancer. Oncogene 33, 1348–1358, doi:10.1038/onc.2013.89 (2013).

32. Nguyen Van Long, F. et al. Low level of Fibrillarin, a ribosome biogenesis factor, is a new independent marker of poor outcome in breast cancer. BMC Cancer 22, doi:10.1186/s12885-022-09552-x (2022).

33. Eeftens, J. M., Kapoor, M., Michieletto, D. & Brangwynne, C. P. Polycomb condensates can promote epigenetic marks but are not required for sustained chromatin compaction. Nature Communications 12, doi:10.1038/s41467-021-26147-5 (2021).

34. Smith, K. S. et al. Bmi-1 Regulation of INK4A-ARF Is a Downstream Requirement for Transformation of Hematopoietic Progenitors by E2a-Pbx1. Molecular Cell 12, 393–400, doi:10.1016/s1097-2765(03)00277-6 (2003).

35. Liang, J. et al. Bmi-1 regulates the migration and invasion of glioma cells through p16. Cell Biology International 39, 283–290, doi:10.1002/cbin.10390 (2014).

36. Xu, J., Li, L., Shi, P., Cui, H. & Yang, L. The Crucial Roles of Bmi-1 in Cancer: Implications in Pathogenesis, Metastasis, Drug Resistance, and Targeted Therapies. International Journal of Molecular Sciences 23, doi:10.3390/ijms23158231 (2022).

37. Wang, M.-C. et al. BMI-1, a promising therapeutic target for human cancer. Oncology Letters 10, 583–588, doi:10.3892/ol.2015.3361 (2015).

38. Chen, D. et al. Targeting BMI1 + Cancer Stem Cells Overcomes Chemoresistance and Inhibits Metastases in Squamous Cell Carcinoma. Cell Stem Cell 20, 621-634.e626, doi:10.1016/j.stem.2017.02.003 (2017).

39. Biehs, B. et al. BMI1 represses Ink4a/Arf and Hox genes to regulate stem cells in the rodent incisor. Nature Cell Biology, doi:10.1038/ncb2766 (2013).

40. Endo, A. et al. Nucleolar structure and function are regulated by the deubiquitylating enzyme USP36. Journal of Cell Science 122, 678–686, doi:10.1242/jcs.044461 (2009).

41. Niu, M.-Y. et al. The Emerging Role of Ubiquitin-Specific Protease 36 (USP36) in Cancer and Beyond. Biomolecules 14, doi:10.3390/biom14050572 (2024).

42. Sun, X.-X., Sears, R. C. & Dai, M.-S. Deubiquitinating c-Myc: USP36 steps up in the nucleolus. Cell Cycle 14, 3786–3793, doi:10.1080/15384101.2015.1093713 (2015).

43. Zhu, S. et al. USP36-Mediated Deubiquitination of DOCK4 Contributes to the Diabetic Renal Tubular Epithelial Cell Injury via Wnt/β-Catenin Signaling Pathway. Front Cell Dev Biol 9, 638477, doi:10.3389/fcell.2021.638477 (2021).

44. Papalazarou, V. & Maddocks, O. D. K. Supply and demand: Cellular nutrient uptake and exchange in cancer. Molecular Cell 81, 3731–3748, doi:10.1016/j.molcel.2021.08.026 (2021).

45. Bulmus Tüccar, T. & Acar Tek, N. Determining the factors affecting energy metabolism and energy requirement in cancer patients. Journal of Research in Medical Sciences 26, doi:10.4103/jrms.JRMS_844_20 (2021).

46. Zhu, S. et al. BMI1 regulates androgen receptor in prostate cancer independently of the polycomb repressive complex 1. Nature Communications 9, doi:10.1038/s41467-018-02863-3 (2018).

47. Singh, S. K., Venugopal, C., Adile, A. A. & Bakhshinyan, D. Bmi1 – A Path to Targeting Cancer Stem Cells. European Oncology & Haematology 13, doi:10.17925/eoh.2017.13.02.147 (2017).

48. Bouffard, S. et al. Fibrillarin is essential for S-phase progression and neuronal differentiation in zebrafish dorsal midbrain and retina. Developmental Biology 437, 1–16, doi:10.1016/j.ydbio.2018.02.006 (2018).

49. Richardson, Lauren A. et al. A Conserved Deubiquitinating Enzyme Controls Cell Growth by Regulating RNA Polymerase I Stability. Cell Reports 2, 372–385, doi:10.1016/j.celrep.2012.07.009 (2012).

50. Fitieh, A., Locke, A. J., Motamedi, M. & Ismail, I. H. The Role of Polycomb Group Protein BMI1 in DNA Repair and Genomic Stability. International Journal of Molecular Sciences 22, doi:10.3390/ijms22062976 (2021).

